# mRNA and Protein Expression of Fetal Insulin Receptor in Breast Cancer Cell Lines

**DOI:** 10.64898/2025.12.15.694420

**Authors:** Xihong Zhang, Albert Barrios, LeeAnn Higgins, Todd Markowski, Kevin Murray, Bruce Witthuhn, Tzu-Yi Yang, Douglas Yee

## Abstract

The insulin receptor (IR) is expressed in breast cancer cells and plays a role in regulating tumor biology. There are two IR isoforms generated from the same gene. Alternate splicing with exclusion or inclusion of exon 11accounts for the two isoforms. The exon 11 excluded isoform (IR-A) is expressed during fetal development while the full-length adult IR (IR-B) is the primary form expressed during adult life. This splice variant results in a 12 amino acid variation in peptide sequence. Breast cancer cells overexpress IR-A with an increased IR-A:IR-B ratio. Most of these data were obtained by examining mRNA expressions. In this work, we examined over 40 breast cancer cell lines and patient tumor samples for mRNA expression of the IR isoforms to show that most cells overexpressed IR-A compared to IR-B. Further we used mass spectrometry to demonstrate IR-A protein expression in the Du4475 cell line which has a high level of IR-A mRNA expression. To our knowledge, this is the first demonstration of IR-A protein expression. Thus, IR-A mRNA and protein expression demonstrate a potential role for this insulin receptor isoform in breast cancer biology.

## Introduction

Elucidating the molecular subtypes of breast cancer has led to novel targeted therapies and improved survival for breast cancer (1). The expression of the estrogen receptor (ER), progesterone receptor (PR), and human epidermal growth factor 2 proto-oncogene (HER2) classify breast cancers into specific subtypes and provide opportunities to individualize therapy based on expression of these key genes. Further mRNA transcriptomic profiling combined with clinical data shows that most patients with ER expressing tumors receive no benefit from cytotoxic chemotherapy (2). Thus, identification of key signaling molecules and translating those findings into clinical practice has resulted in the observed improved outcomes.

The IR and IGF-1R have long been reported to be overexpressed in breast cancer and to promote tumor growth (3,4). IR and IGF-1R are both transmembrane tyrosine kinase heterotetrametric proteins, where the two extracellular *α*-subunits is the site of insulin binding and two transmembrane β-subunits, the site of tyrosine kinase activity, initiate the signaling cascade for growth (5,6). IR and IGF-1R differ in their interactions and functions, but they are structurally homologous. Hence, hybrid receptors, IR/IGF-1R, exist and add more to the complexity to the IR/IGF system (7). The insulin receptor was reported to associate with increased tumor size and weight, and increased migration and proliferation, by activating ERK via phosphorylation on MCF7 cells (8). In addition, it was reported that IGF-1R knockdown resulted in no changes to growth and identified that IR send mitogenic signals during insulin or insulin-like growth factor 2 (IGF-2) binding (9). Insulin receptor was also implicated in stem-like cell phenotype, angiogenesis, and other aggressive tumor behaviors in triple negative breast cancer via induction of *vegfa, pdgfa*, and *serpine2* (10). Thus, both IGF-1R and IR could play a role in cancer biology.

Two isoforms of the insulin receptor exist. The human insulin receptor gene is composed of 22-exons spanning 120 kb in chromosome 19 (11). Alternate splicing results of exon 11 results in the deletion of 12 amino acids from the fetal form of the insulin receptor (IR-A) while the adult form (IR-B) maintains this exon. Both isoforms respond to insulin, but IR-A has a higher affinity for IGF-II than IR-B. As IGF-II has been implicated in cancer, the quantification of the isoforms may have therapeutic implications. Further, in endocrine resistant breast cancer, IR-A is expressed at higher levels than IR-B or IGF-1R and could be a therapeutic target.

Techniques to detect and quantify mRNA expression of IR isoforms have been described (12) and suggest that cancer cells express IR-A at higher levels than IR-B. To date, there have been no reports of the detection of IR-A protein given the very close homology to IR-B. In this study, we evaluated mRNA and protein expression of IR-A. We found that IR-A mRNA expression in breast cancer cell lines is proportional to protein expression detected by mass spectrometry.

## Material and Methods

### Human breast cancer cells, tissues, and reagents

The ATCC Breast Cancer Cell Panel was purchased from ATCC (ATCC 30-4500K) and cultured according to ATCC site instructions. MCF-7L cells were a gift from C. Kent Osborne (Baylor College of Medicine, Houston, TX) and maintained in improved MEM Richter’s modification medium (zinc option) supplemented with 11.25 nmol/L insulin, 1% penicillin and streptomycin, and 5% fetal bovine serum.

The heavy-labeled (C13- and N15-containing) IR-A peptide was from Thermo Scientific with the sequence TFEDYLHNVVFVPRPS. IR, IR-A and IR-B primer sequences were as described (12) and synthesized Integrated DNA Technologies. IR antibody was purchased from Santa Cruz Biotechnology (sc-57342) and secondary goat anti mouse IgG (H+L)-HRP conjugate antibody for immune blots was from BioRad (Cat # 1706516).

Normal human tissues were received obtained from the UMN cohort (IRB Approval Study# 1409E53504). They were collected from archived pathology tissue blocks with de-identified clinical data; all participants had agreed to the institution’s standard consent for research participation. The breast cancer samples were collected as previously described (13).

### Reverse transcription-quantitative real-time polymerase chain reaction (RT-qPCR)

Total RNA was made from cells or tissue by using TriPure Isolation Reagent according to the manufacturer (Roche Diagnostics, Indianapolis, IN). A total of 2 μg of RNA was reverse transcribed using the Transcriptor first strand cDNA synthesis kit (Roche), and quantitative PCR was performed using the FastStart universal SYBR Green kit according to the manufacturer’s recommended protocol (Roche). When prepping cDNA samples for quantitative PCR, IR-specific primers for IR-A, IR-B, IGF-1R, and total IR were utilized (12). Data was collected and analyzed using Bio-Rad qPCR machine and software (Ribosomal Protein Lateral Stalk Subunit P0 – RPL0 was used as reference gene).

### Membrane preparation and immunoprecipitation

Cultured cells were washed 2x with cold PBS. Cell pellet was suspended in 1 ml ice cold membrane lysis buffer (20mM Tris HCL pH 7.4, 2 mM MgCl2, 1 mM EDTA and Protease inhibitor cocktail from Roche Cat# 11836153001). Cells were sheared through a 26G needle. Sheared cells were centrifuged at 1000rpm for 5 minutes at 4°C and pellet was discarded. The suspension was recentrifuged at 15000 rpm for 30 minutes in the cold. The pellet, primarily cell membranes, was resuspended in TNESV buffer and recentrifuged at 13750 rpm for 20 minutes in the cold. 500 μg supernatant protein was utilized for immunoprecipitation as previously described (14). Briefly 1:50 IR antibody in a volume of 0.5 ml protein extract was incubated for 2 h at 4°C. Sample were processed with gentle agitation at 4°C overnight with 25μl protein A/G plus agarose beads. Beads were washed five times and pulldown proteins were used for immunoblotting with IR primary and secondary antibodies.

Excised gel regions were subjected to in-gel digestion as described previously (15) with the following differences. The proteolytic enzyme used for the in-gel digestion was LysC (Promega) and the alkylation reagent was 55 mM iodoacetamide. After overnight incubation, 5 μl of 100 fmoles/μl heavy peptide from protein of interest was spiked into the digestion. Gel bands were extracted and dried down in a vacuum centrifuge. Each sample was cleaned with a C18-like Stage tip (16). Eluates were vacuum dried.

### Detection of IR-A via Orbitrap Fusion LC-Mass Spectrometry

We reconstituted the dried LysC peptide mixtures in 98:2:0.1,H2O:acetonitrile (ACN):formic acid (FA) and analyzed ∼200 nanogram of each sample by capillary LC-MS on an Orbitrap Fusion mass spectrometer (Thermo Fisher Scientific, Inc., Waltham, MA) online with a Thermo UltiMate™ 3000 RSLCnano LC system. Peptides were separated on a 75 cm self-packed C18 capillary column with 100 um inner diameter, with Dr. Maisch GmbH ReproSil-PUR 12 Å C18-AQ, 1.9 um particle size; the column was maintained at 55°C with a column heater from Sonation (Biberach, Germany). Peptides were loaded directly on column at 325 nl/min with 98:2:0.1,H2O:ACN:FA. We performed gradient elution of peptides with 5 – 35% solvent B (0.1% FA in ACN) in 70 minutes, 35 – 90% B from 77 – 83 min, and held at 90% B for 6 min. Solvent A was 0.1% FA in H2O, solvent B was 0.1% FA in acetonitrile and the column flowrate was 325 nl/min. We operated the mass spectrometer with the following parameters: ESI voltage 2.1kV, ion transfer tube 275 °C; Orbitrap MS1 scan 120k resolution in profile from 360 – 1580 m/z with 50 msec injection time, 125% normalized automatic gain control (AGC); the number of dependent scans was set to the top 20. Tandem MS parameters were: precursor isolation width 1.6 Da, collision induced dissociation (CID) activation with 35% collision energy, 10 ms activation time and 0.25 activation Q, ion trap detection, 35 msec injection time, 1x 10^4^ automatic gain control (AGC), dynamic exclusion duration 20 sec with +/-10 ppm mass tolerance.

### PEAKS Studio Database Search

We used Peaks® Studio v10 (PMID: 14558135 Bioinformatics Solutions, Inc, Waterloo, ON CA) for interpretation of tandem MS and protein inference. Search parameters were: human universal proteome sequence database (UP000005640) from UniProt.org (July 12, 2019) merged with the short form of human insulin receptor (Uniprot P06213-2) and common lab contaminant proteins from http://www.thegpm.org/crap/; parent mass error tolerance 20.0 ppm; fragment mass error tolerance 0.6 Da; precursor mass search type monoisotopic; enzyme trypsin with maximum 2 missed cleavage sites and semi-specific digest mode; variable modifications methionine oxidation, (+15.9949) pyroglutamic acid from glutamine (-17.0265), deamidation of asparagine and glutamine (+0.9840), protein N-terminal acetylation (+42.0106); fixed modification carbamidomethyl (CAM) cysteine (+57.0215); maximum variable modifications per peptide 3; false discovery rate calculation On; spectra merge option OFF, charge correction off, charge filter 2 – 6; the de novo sequencing parameters were: parent mass tolerance 20 ppm, fragment mass tolerance 0.6 Da, fixed modification CAM cysteine and variable modification oxidized methionine. The peptide and protein lists were filtered at 1% false discovery rate.

### LC-MS/MS Parallel Reaction Monitoring (PRM) Analysis of IRA

We analyzed the LysC peptide mixtures identically to the discovery experiment except for the following LC and MS parameters. We analyzed approximately 200 ng of peptide mixture with 100 femtomole of the heavy peptide. We performed gradient elution of peptides with 5 – 60% solvent B (0.1% FA in ACN) in 55 minutes, 60 – 90% B from 55 – 60 min, and held at 90% B for 8 min. Solvent A was 0.1% FA in H2O, solvent B was 0.1% FA in acetonitrile, and the column flowrate was 325 nl/min. The mass spectrometer parameters were: Orbitrap MS1 range 380 – 1580 m/z, 100% AGC. Tandem MS parameters were: high-energy collisional dissociation (HCD) activation precursor isolation width 1.6 Da, with 30% collision energy, orbitrap detection at 30k resolution (at 200 m/z), 54 msec injection time, 5x 10E4 automatic gain control (AGC). We monitored the doubly and triply charged ions for light (640.6599 and 960.4862 m/z) and heavy-labeled peptide (643.9959 and 965.4903 m/z) in PRM mode.

We processed the PRM data in Skyline (version 22.2.0.527, Skyline Team from University of Washington, Seattle, WA - https://skyline.ms) with a spectral library exported from PRM data analyzed in PEAKS Studio with the same processing parameters used for the discovery experiment. Peak areas were calculated from transitions of the triply charged ions for the heavy and light peptides. Skyline processing settings included: 0.05 m/z tolerance for the spectral library match, 4 product ions, isotope labels 13C(6) 15N(4) R.

## Results

### Expression of IR isoforms in breast cancer cell lines and normal tissues

Expression of insulin receptor subtypes varied between cancer cells and normal tissues. We studied the breast cancer cell line MCF-7 where we have previously used RNAse protection assays to show that IR-A and IR-B expression were both expressed by these cells (13). Figure 1A shows MCF-7 expressed high levels of IR-A. In comparison to the insulin target tissues (adipose tissue, liver, and muscle) total levels of IR were lower than in the breast cancer cells. Notably, IR-B is the predominant subtype expressed in all adult tissues although muscle had a high level of IR-A mRNA.

**Figure 1.**
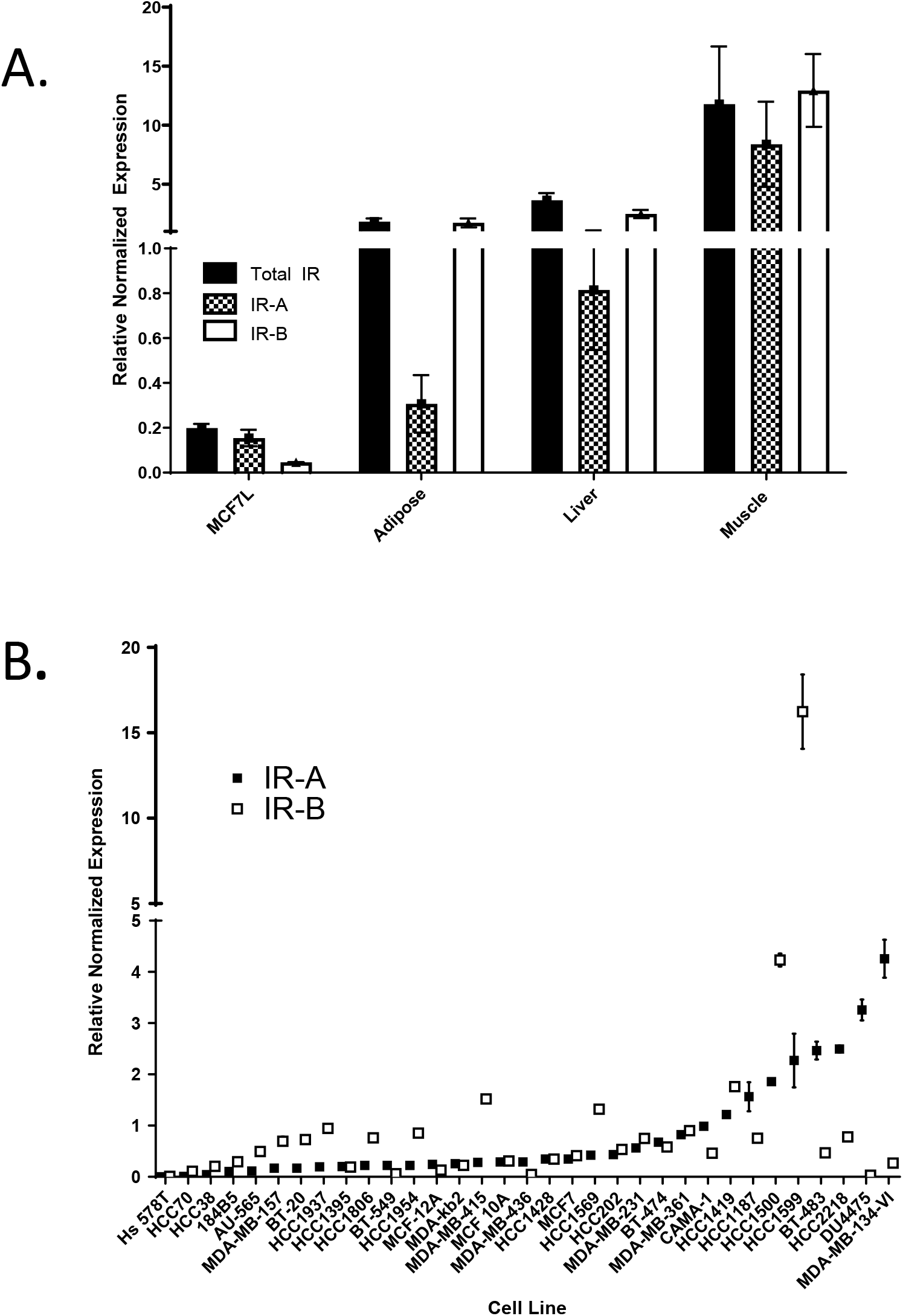
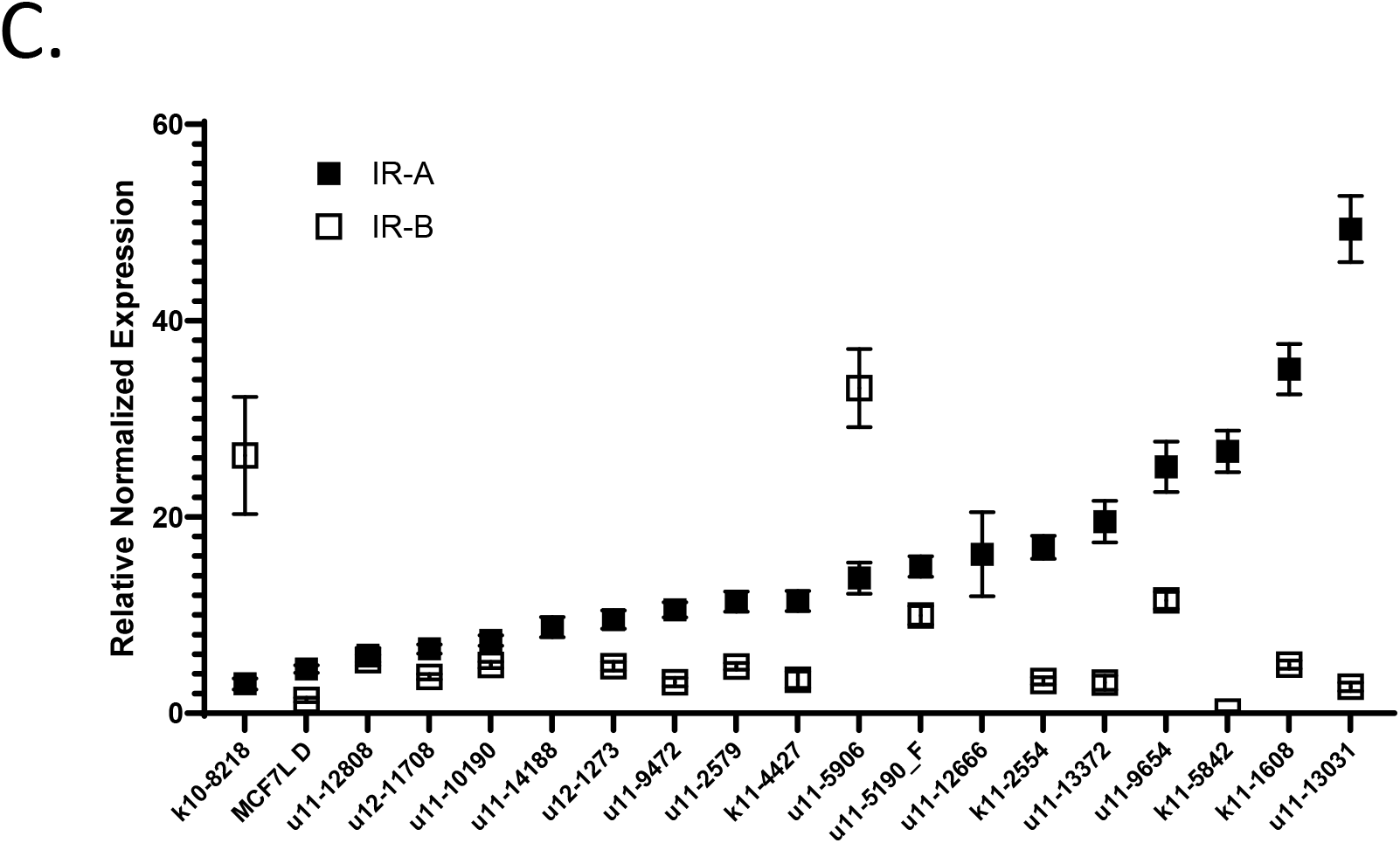
mRNA expression of IR isoforms in breast cancer cell lines, normal tissues, and primary breast cancers. A. Expression of total IR, IR-A, and IR-B mRNA in MCF-7 cells and human tissues B. Expression of IR-A and IR-B mRNA expression in breast cancer cell lines C. Expression of IR-A and IR-B in breast cancer specimens

Figure 1B demonstrates IR-A and IR-B mRNA expression in breast cancer cell lines. Most cells expressed both isoforms and there was heterogeneity of expression in the cell lines. However, some cells expressed higher levels of IR-A over IR-B. When breast cancer tissues were analyzed (Figure 1C) there was a higher level of IR-A mRNA in most specimens compared to IR-B.

### Detection of insulin receptor isoform A by mass spectrometry

To detect IRA peptide, we utilized the mRNA data to select a cell line with high expression of IR-A. We enriched IR protein via immunoprecipitation using Du4475 cell line which had a high level of IR-A mRNA expression (Figure 1B). 500 mg membrane lysate protein was immunoprecipitated with IR antibody in B (Figure 2A). The enriched IR was subjected to SDS-PAGE using 50 µg total lysate protein as control (Figure 2A), then isolated protein at 135 kDa (IR alpha subunit) was excised from the gel and used for mass spectrometry analysis (Figure 2B).

**Figure 2.**
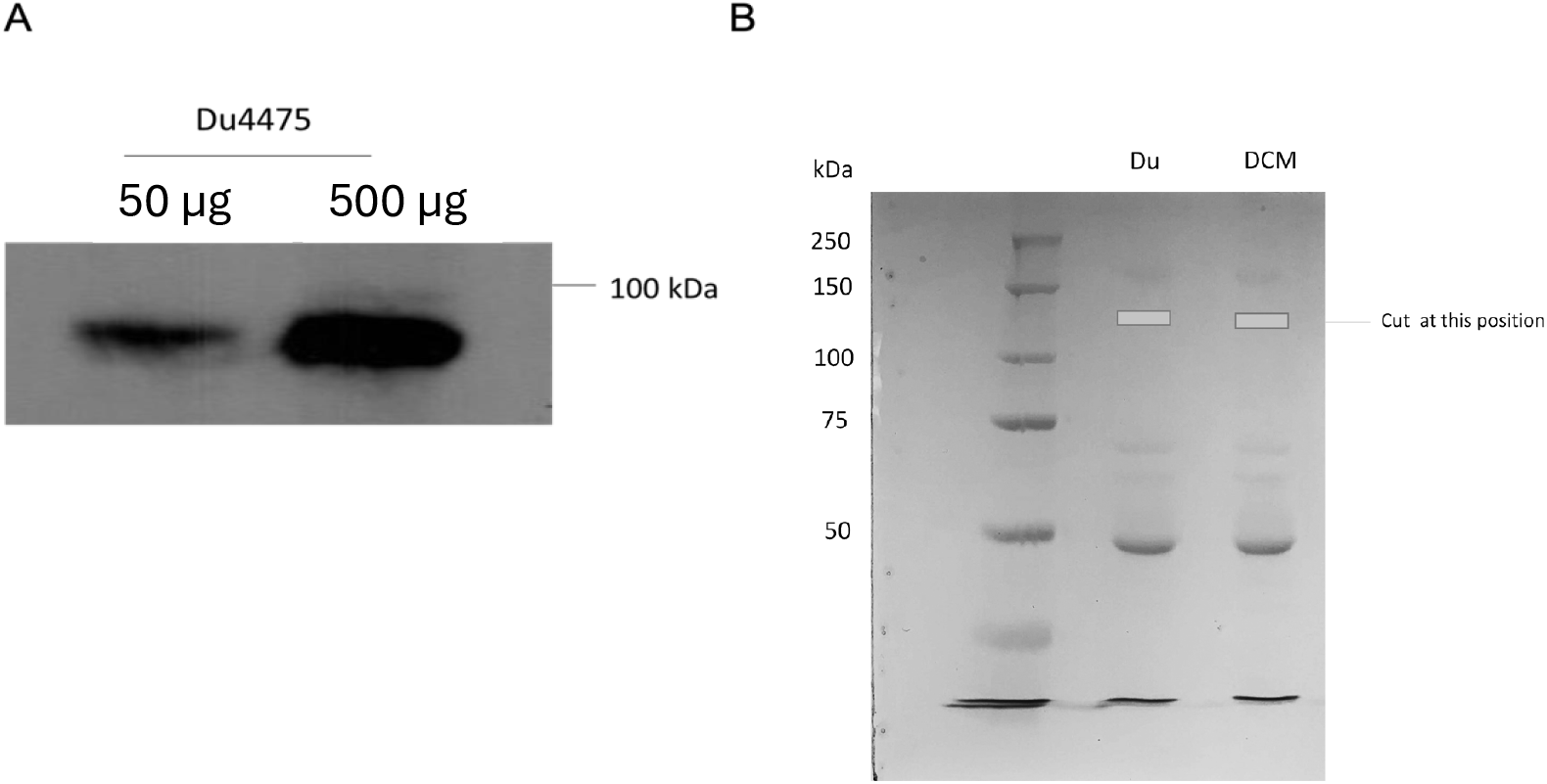
Detection of IR in breast cancer cells. A. Immunoprecipitation of DU4475 membrane protein with IR antibody using 50 and 500µg of lysate B. Immunoprecipitated proteins were excised after PAGE. DU is 100% DU4475. DCM is a 50:50 mixture of DU4475:MCF7L

We performed a mass spectrometry (MS)-based discovery proteomics experiment using endoproteinase Lys-C (Lys-C) and detected IR-A (P06213-2 Uniprot accession) with 20% coverage (by amino acid count) from 36 unique peptides. We detected a semi-specific, Lys-C peptide, TFEDYLHNVVFVPRPS ([M +2H]^2+^, monoisotopic, theoretical m/z = 960.4862), that contains the sequence unique to IR-A [Uniprot accession P06213-2] and provided differentiation from IR-B (P06213-1 Uniprot accession). We compared the tandem MS data for the synthetic, heavy-labeled peptide (13C(6), 15N(4)-arginine) to the experimental data and confidently verified the peptide match (Figure 3A).

**Figure 3.**
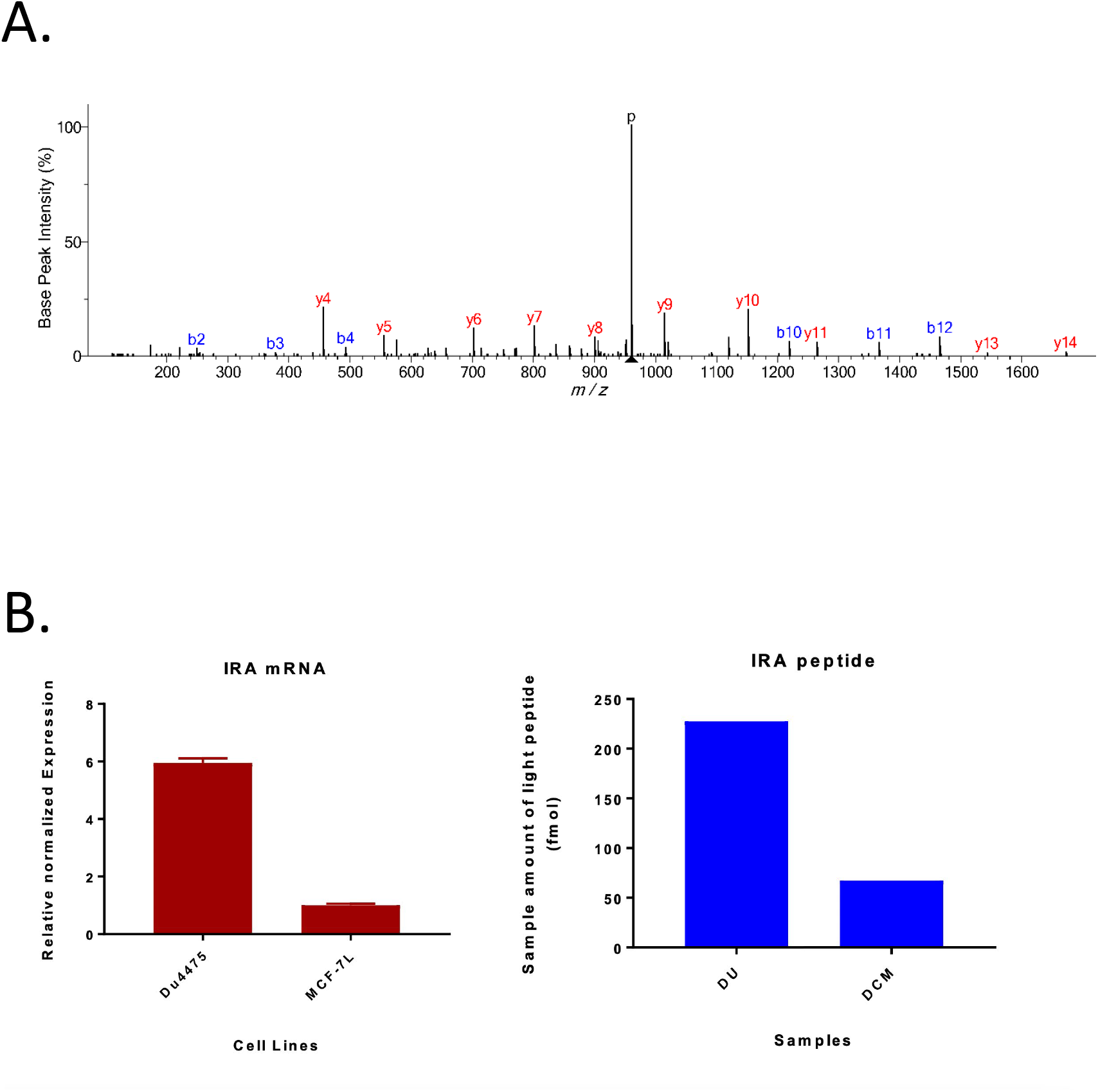
Detection of IRA specific peptide in DU4475 and DCM by mass spectrometry.=. A. DCM membrane proteins recovered from SDS-PAGE gel were digested by using endoproteinase Lys-C. The resulting peptides underwent mass spectrometry analysis. Tandem MS spectrum shown here is from the [M + 2H] +2 precursor 960.4866 m/z (theoretical, monoisotopic mass with 0.5 ppm mass error). The spectrum is annotated with the diagnostic b- and y-type fragment ions. The following software analysis indicates the unique IRA peptide sequence of FEDYLHNVVFVPRPS. B. Correlation of mRNA and protein expression. IR-A mRNA was measured in Du4475 and MCF-7L cells. Peptide expression was measured by mass spectrometry in Du4475 (DU) and DCM (Du4475 to MCF-7L lysate ratio is 1:1). Protein concentrations were determined with heavy peptide as noted in the Materials and Methods. Levels were proportional between mRNA expression (left panel) and peptide detection (right panel).

For a quantitative measurement, we conducted a parallel reaction monitoring (PRM) experiment with heavy-labeled peptide spiked into the peptide mixtures immediately after in-gel digestion with Lys-C. We report the relative levels of IR-A protein in Du4475 (Du) and equal mixture of Du4475+MCF7 (DCM) cells using the heavy-labeled peptide as the internal standard. The peak areas of 4 transitions for the light peptide were summed and divided by the same peak areas for the internal standard. We showed that IRA was 3.4x higher in abundance in Du4475 cells compared to Du4475+MCF7 (228 fmol of peptide TFEDYLHNVVFVPRPS was measured in Du versus 67 fmol of peptide in DCM (Figure 3B).

## Discussion

The roles of the IGF signaling system have been well studied and reported extensively (17). There has been evidence that the IR-A isoform is more highly expressed in cancer cells and tissues (18), but these observations have been largely based on mRNA expression. Different biological effects have been attributed to the IR isoforms (6,11), thus understanding the expression in cells may have physiologic and pathophysiologic consequences. However, the demonstration of IR-A protein expression has been difficult given the small difference in the protein expression between the two isoforms; the IR-A isoform is missing exon 11 which results in a deletion of 12 amino acids when compared to the adult IR-B isoform.

Here we confirmed that IR-A mRNA is detected in breast cancer cell lines and tissues. While both receptors were identified, there was a predominance of IR-A mRNA expression compared to IR-B. Using these findings, we used mass spectrometry to identify the peptide sequence for IR-A. Since exon 11 is not included in IR-A, we created a peptide that was predicted by the direct splicing of exon 10 to exon 12 of the IR gene. A stable isotopic peptide was synthesized representing the unique LysC peptide sequence corresponding to the direct splicing of exon 10 to exon 12 of the IR gene. This peptide corresponded to the predicted amino acid sequence of IR-A. The heavy LysC peptide was used to optimize mass spectrometry detection, and the heavy peptide was used to identify the correct IR-A peptide from unlabeled cell lysates. Using this strategy, we were able to demonstrate the presence of the peptide in a cell line with high expression of IR-A. Further, when we mixed lysates from a high and low expressing cells, the detection of IR-A was proportional to the mRNA expression. To our knowledge, this is the first demonstration of the presence of IR-A protein in human cells.

In this work, we could detect the appropriate IR-B protein sequence by mass spectrometry (data not shown) but could not quantify the expression of IR-B protein in these using the same technique as we did for IR-A. The IR-B heavy peptide formed a secondary structure around the prolines in the IR-B peptide that are known to affect chromatography(19). Ideally, it would be instructive to understand if the mRNA and protein ratios of expression were matched. However, the IR-B peptide is hydrophilic in proximity to the splice site. We surmise that this difference promotes a secondary structure in a proline-rich sequence of the IR-B peptide affecting its ability to be detected by mass spectrometry. Further work will be needed to accurately quantify IR-B protein.

In conclusion, we were able to use mass spectrometry to validate protein expression of IR-A expression in cells that express high levels of IR-A mRNA. Further work on developing techniques to distinguish between the IR isoforms could yield new information on breast cancer biology and develop new biomarker-driven targeted therapies for IR-A expressing tumors.

